# Benchmarking of lightweight-mapping based single-cell RNA-seq pre-processing

**DOI:** 10.1101/2021.01.25.428188

**Authors:** A. Sina Booeshaghi, Lior Pachter

## Abstract

We compare and benchmark the two lightweight-mapping tools that have been developed for pre-processing single-cell RNA-seq data, namely the kallisto-bustools and Salmon-Alevin-fry programs. We find that they output similar results, and to the extent that there are differences, they are irrelevant for downstream analysis. However, the Salmon-Alevin-fry program is significantly slower and requires much more memory to run, making it much more expensive to process large datasets limiting its use to larger servers.

## Introduction

The analysis of single-cell RNA-seq data begins with pre-processing of sequenced reads to generate count matrices. This analysis prerequisite is the analog of quantification of bulk RNA-seq data, however it is considerably more complex (Luecken and Theis 2019). Several different technology-specific approaches to single-cell pre-processing have been implemented as part of proof-of-principle demonstrations of the experimental techniques, e.g. (Cao et al. 2018; Macosko et al. 2015), but such custom solutions do not scale well and as datasets grow in size (Svensson, da Veiga Beltrame, and Pachter 2020; Cao et al. 2020) algorithmically sound and computationally efficient tools are needed.

One promising approach to single-cell RNA-seq pre-processing is lightweight-mapping, which was initially developed for bulk RNA-seq in the form of pseudoalignment implemented in the kallisto program (Bray et al. 2016), and later re-implemented in the Salmon program (Patro et al. 2017). More recently, the Salmon pseudoalignment algorithm has been modified to perform “selective alignment” (Srivastava et al. 2020) and a sketch alignment strategy was recently implemented in RainDrop, though the tool is limited to processing only a single data type and for that reason we do not consider it here (Niebler et al. 2020). Following the naming convention of (Srivastava et al. 2020) we collectively refer to pseudoalignment and selective alignment as “lightweight-mapping” procedures to distinguish them from standard alignment algorithms. The two bulk RNA-seq lightweight-mapping based quantification programs have already been adapted for single-cell RNA-seq: the Alevin tool (Srivastava et al. 2019) adapts the Salmon program to make it suitable for single-cell RNA-seq pre-processing, and the bustools suite of programs (Melsted et al. 2019) can be used to assemble single-cell RNA-seq pre-processing workflows using a novel format (Melsted, Ntranos, and Pachter 2019) that can be generated, in principle, by any lightweight mapping program. The kallisto-bustools program has been shown to be much faster and more efficient than other tools for single-cell RNA-seq pre-processing (Srivastava et al. 2019; Melsted et al. 2019; Wu and Zhang 2020) and we expect that as single-cell RNA-seq datasets scale to millions of cells and tens of billions of reads (Cao et al. 2020), this performance advantage will become essential for the generation of transparent and readily reproducible results.

A key difference between kallisto-bustools and Salmon-Alevin is that unlike Salmon-Alevin, the kallisto-bustools approach is centered on the generation and manipulation of BUS (Barcode, UMI, Set) files (Melsted, Ntranos, and Pachter 2019). This serves to decouple the technology dependent pseudoalignment step from the other single-cell RNA-seq pre-processing steps such as barcode error correction, unique molecular identifier collapsing and read counting. This modular structure makes it easy to pre-process data from different technologies in a uniform way, and enables a large number of downstream analyses by virtue of easy customization of individual pre-processing steps. Alevin, on the other hand, is integrated into the Salmon software, and bundles a solution to all of these tasks into a single program. This has two disadvantages: it makes it harder to customize workflows for different technologies and distinct pre-processing requirements, and it makes it harder to substitute algorithmic components; any change to an algorithm requires familiarity with and modification of a large, complex, and tightly integrated codebase.

In light of the modular kallisto-bustools solution to single-cell RNA-seq pre-processing and the advantages it confers over Alevin, the authors of Alevin have recently adapted Salmon-Alevin to output a BUS-like file, and have reimplemented the bustools programs, with minor modifications. Their reimplementation is called Alevin-fry (Sarkar et al. 2020). Despite being written in different programming languages (bustools in C++ and Alevin-fry in Rust), the two solutions for single-cell RNA-seq pre-processing are now identical in structure and have the same functionality, raising the question of whether there are substantial differences in usability, output, and efficiency. To answer this question, we have performed a comprehensive comparison and benchmarking of kallisto-bustools and Salmon-Alevin-fry, taking particular care to avoid the self-assessment trap (Teng et al. 2016; Buchka et al. 2021). To do so we adhere carefully to developer recommendations, and compare the programs with respect to a large number of datasets that include data from a wide variety of organisms and tissues. We have also included code reproducing our results, so that users can explore the tools and data for themselves, without having to take our word about performance, features, and benchmarks.

## Results

The BUS format was first published on November 21, 2018 (Melsted, Ntranos, and Pachter 2019, 2018) followed by a release of the open-source kallisto-bustools software for single-cell RNA-seq pre-processing on December 5, 2018 and a publication on June 17, 2019 (Melsted et al. 2019). For the comparisons performed here, we used the latest versions of kallisto (0.46.2) and bustools (0.40.0). In order to compare results generated with kallisto-bustools to the Salmon-Alevin-fry reimplementation published on November 5, 2020 (Sarkar et al. 2020) and released on December 2, 2020, we ran the latest releases of Salmon-Alevin (1.4.0) and Alevin-fry (0.1.0). The components of the programs were benchmarked individually, and corresponding parts were compared to each other directly. The kallisto-bustools approach is based on the BUS (Barcode, UMI, Set) format, which was introduced in (Melsted, Ntranos, and Pachter 2019) as a crucial data type and format for decoupling pseudoalignment from the downstream steps of pre-processing. The Salmon-Alevin-fry pipeline appears to use a similar format called RAD (Reduced Alignment Data) that serves the same purpose, although there is no official specification for the format thus tying it to the Alevin-fry software. The bustools programs used to produce a complete pre-processing workflow by manipulating BUS files are mirrored in the Alevin-fry tools that reimplement them (Figure 1).

**Figure 1:**
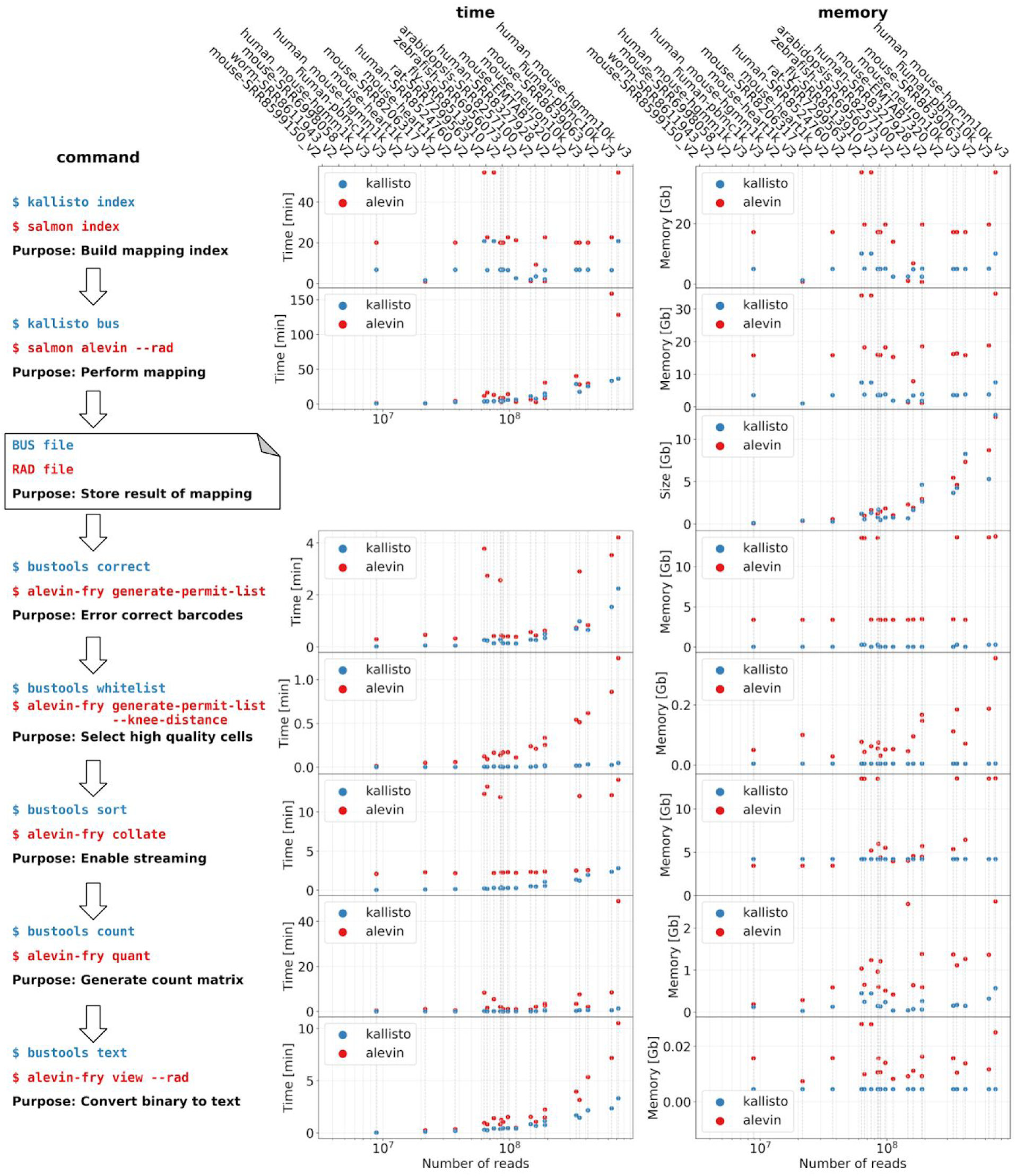
The corresponding components of the kallisto-bustools and Salmon-Alevin-fry workflows (left). After constructing an index and performing lightweight-mapping of the reads, the output is saved to a file. Subsequently this file serves as a substrate for performing barcode error correction and UMI counting among other tasks; time to run each command for the 20 datasets analyzed using 10 threads (middle); memory requirements (right). [[Code](https://github.com/pachterlab/BP_2021/blob/main/analysis/notebooks/memtime.ipynb)]

To ensure that the programs were run according to developer recommendations, we examined the documentation and ran the programs accordingly. On the Salmon documentation page the authors state that “When salmon is run with selective alignment, it adopts a considerably more sensitive scheme that we have developed for finding the potential mapping loci of a read” (Patro et al. 2021b) so we ran Salmon in selective alignment mode. The Alevin documentation (Patro et al. 2021a) states that “we recommend using ~10 threads which in our testing gave the best memory-time trade-off”. We adopted this recommendation. To prepare the index, we followed the recommendation of “using selective alignment with a decoy-aware transcriptome”, and constructed a full-genome “decoy-aware” transcriptome index based on the recommendation that “this scheme provides a more comprehensive set of decoys” (Patro et al. 2021b) and instructions on the Alevin-Tutorial page (COMBINE-lab n.d.). For the purposes of ensuring fair runtime and memory benchmarks we ran kallisto and bustools with the same number (10) of threads as allocated to Salmon-Alevin-fry, and otherwise ran all commands without setting parameters so that default values would be used, with the exception of the -m flag for bustools limiting it to 4Gb of RAM. We note that in the case of Alevin, output logs from the program left us uncertain as to what those default values were (Supplementary Figure 1) and lack of documentation for some parameters did not allow us to resolve these ambiguities. For example, the output log for the mouse-EMTAB7320_v2 dataset reported the `minScoreFraction` parameter as set to both 0.65 and 0.87. Additionally, the log describes the `consensusSlack` parameter as set to a default of both 0.35 and 0.6.

To compare the results of running the programs we assessed them on 20 datasets, including published single-cell RNA-seq datasets from *H. sapiens* (Merino et al. 2019; Carosso et al. 2018), *M. musculus* (Miller et al. 2019; O’Koren et al. 2019; Jin, Warunek, and Wohlfert 2018; Delile et al. 2019; Guo et al. 2019), *R. norvegicus* (Mays et al. 2018), *C. elegans* (Packer et al. 2019), *A. thalania* (Ryu et al. 2019), *D. melanogaster* (Mahadevaraju et al. 2020), *and D. rerio* (Farrell et al. 2018), as well as three datasets with cells from both *H. sapiens* and *M. musculus* distributed by 10X Genomics (“Datasets -Single Cell Gene Expression -Official 10x Genomics Support” n.d.), and several other 10X Genomics distributed datasets. For each dataset, we performed a detailed comparison of the output of kallisto-bustools to Salmon-Alevin-fry (Figure 1, Supplementary Figure 2), to assess the extent to which downstream analyses would be affected by choice of program. The dataset for Figure 1 was chosen from among the 20 datasets because it is the dataset benchmarked in (Sarkar et al. 2020).

In analyzing the PBMC 10k cell dataset from 10X Genomics (Figure 1), we first examined the “knee plot” (Figure 1a), which is used to filter high quality cells to be used for downstream analysis. In order to enable a direct comparison, we examined the similarity of the knee plots for cells deemed high quality when filtered by both the whitelists produced by Salmon-Alevin-fry (‘alevin-fry generate-permit-list --knee-distance’), kallisto-bustools (‘bustools whitelist’) and Cell Ranger (Zheng et al. 2017). We found that the knee plots were nearly identical for the PBMC 10k dataset and for the other 19 datasets analyzed (panel A subplots, Supplementary Figure 2). Next, we examined the total counts per cell, which provides a “pseudo-bulk” comparison of the datasets (Figure 1B). Again, the results were nearly identical (kallisto had slightly more counts per cell by a small margin), which means that normalization procedures utilizing counts per cell would yield nearly identical results when applied to the data.

The genes detected plot (Figure 1C) shows the number of genes detected in each cell as a function of unique molecular identifiers (UMIs) in that cell. With the exception of cells with very few counts, which are filtered using the knee plot, the results were nearly identical. To the extent that there were any differences visible between the Salmon-Alevin-fry and kallisto-bustools cells in the plots in Figures 2 panels A-D, we noticed the cells were of poor quality with high mitochondrial content (Supplementary Figure 3). To assess the similarity in gene counts, we computed the correlation between gene counts for each cell (Figure 1D). Most of the correlations of high quality cells in the comparison between kallisto-bustools and Salmon-Alevin-fry are above 0.9 in the datasets we examined, and in many cases exceed 0.95. This is very similar to those between kallisto-bustools and Cell Ranger (Melsted et al. 2019).

**Figure 2:**
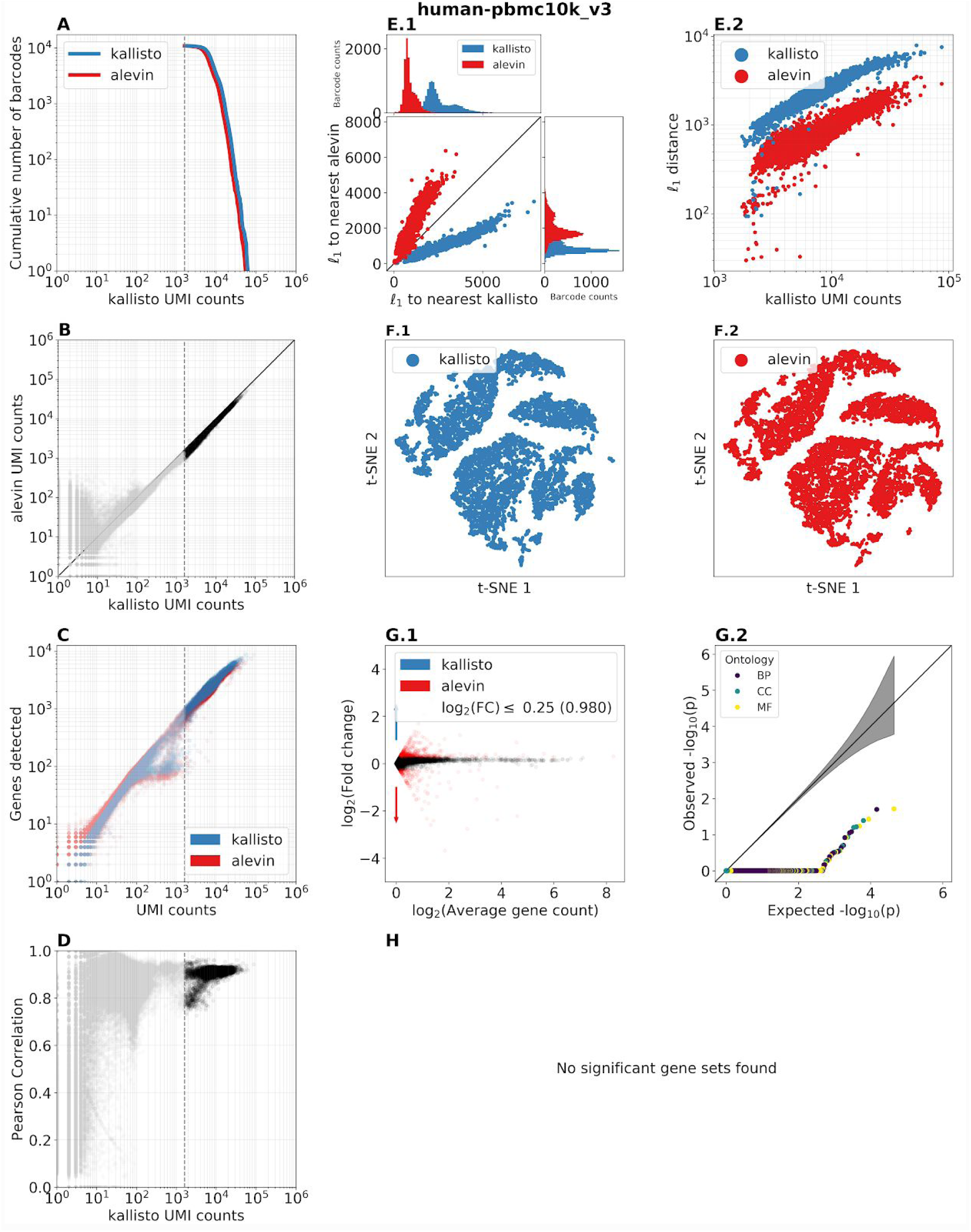
Benchmarking of the 10x Genomics pbmc10k_v3 dataset. Panels A-H display concordance with respect to cell and gene counts. Panels E and F show that distances between cells are preserved. Panels G and H show that no gene sets are significantly perturbed. [[Code](https://github.com/pachterlab/BP_2021/blob/main/analysis/scripts/mkplot.py)]

Two frequently practiced steps in single-cell RNA-seq analysis are clustering of cells and dimensionality reduction of the data. The relevant information for these are the distances between cells. To examine whether clustering algorithms and dimensionality reduction methods would perform differently with kallisto-bustools versus Salmon-Alevin-fry counts, we therefore examined the distances between cells (Figure 1E). For each kallisto-bustools cell, we compared its *l*_1_ distance to the nearest Salmon-Alevin-fry cell, and also its distance to the nearest different kallisto-bustools cell. Figure E.1. shows that kallisto-bustools cells are much closer to their doppelgänger Salmon-Alevin-fry cells, than to other kallisto-bustools cells, and similarly Salmon-Alevin-fry cells are much closer to their doppelgänger kallisto-bustools cells, than to other Salmon-Alevin-fry cells. Figure E.2. shows that while such distances grow with the number of UMI counts per cell, the relative differences between doppelganger cells, vs. distinct cells remains the same. The close distance between kallisto-bustools cells and their doppelgänger Salmon-Alevin-fry cells is the reason t-SNE visualizations based on the two programs are highly similar (Figure 1F).

Finally, to assess whether any specific genes or gene categories are counted differently by kallisto-bustools or Salmon-Alevin-fry, we produced an MA plot (Figure 1G.1) and examined whether specific gene categories were quantified different (Figures 1.G.2 and 1.H). The MA plot shows that not only are fold-change differences in gene counts tiny (98% of genes were within 1.2 fold-change), there are hardly any genes with large fold change differences that are of high abundance. Additionally, there were no gene sets significantly perturbed. In the other 19 datasets analyzed (Supplementary Figure 2), there were sometimes a few gene sets that were perturbed, but these consisted almost entirely of genes encoding for ribosomal proteins and other classes of genes that are typically filtered out prior to analysis of single-cell RNA-seq data. Moreover, the number of genes in these classes constituted a tiny fraction of the genes in the genome. We note that in a handful of datasets there was some divergence in counts observed in poor quality cells, likely the result of kallisto-bustools and Salmon-Alevin-fry treating ambiguously mapped reads differently: Salmon-Alevin-fry by default disambiguates reads with an expectation-maximization algorithm, whereas kallisto-bustools by default does not (though the option exists in ‘bustools count’).

The total run times per dataset of the Salmon-Alevin-fry programs were consistently much higher than that of kallisto-bustools, and the memory requirements were much higher as well (Figure 3). Salmon-Alevin-fry was on average 3 times slower than kallisto-bustools, a result consistent with the benchmarks in (Li et al. 2020). Importantly, while kallisto-bustools ran in under 4Gb of RAM for all datasets except the human-mouse mixed samples, Salmon-Alevin-fry required up to 18.8 Gb of RAM for some of those datasets. For the human-mouse mixed samples kallisto-bustools required 7.5 Gb and Salmon-Alevin-fry required 34.7 Gb.

**Figure 3:**
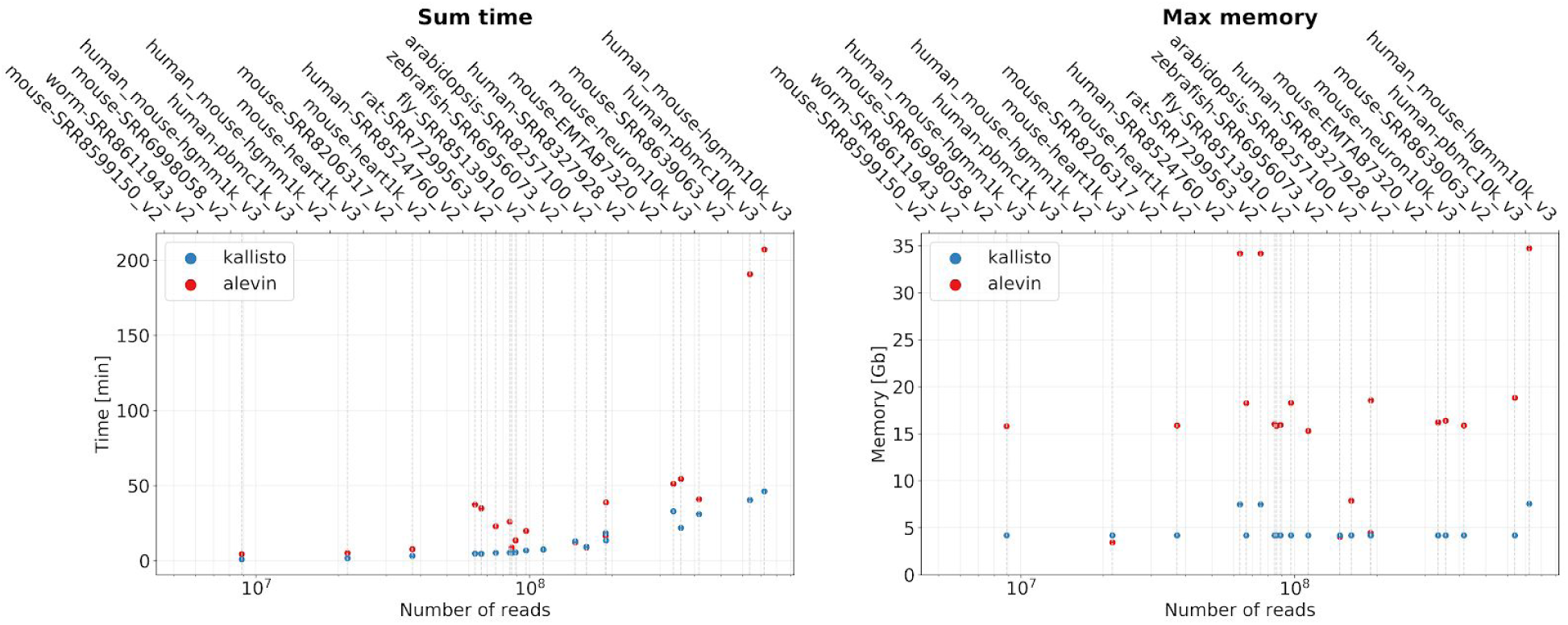
Total run times for each of the datasets with Salmon-Alevin-fry and kallisto-bustools (left); memory needed to run each of the datasets (right). [[Code](https://github.com/pachterlab/BP_2021/blob/main/analysis/notebooks/memtime.ipynb)]

## Discussion

Our benchmarks of kallisto-bustools and Salmon-Alevin-fry reveal that Salmon-Alevin-fry is slower than kallisto-bustools and uses much more memory, yet the results it outputs are nearly identical to those of kallisto. This finding is in contradiction to the claims of some of the Salmon authors, e.g. “Alevin…is among the fastest and most accurate ways to process many types of single-cell protocols to produce expression estimates” (Caulfield 2020). The inefficiency of Salmon-Alevin-fry has consequences in terms of compute costs. We priced the cost of running kallisto-bustools and Salmon-Alevin-fry on the datasets used in this paper, using pricing from Amazon Web Services. We found that for the PBMC 10k dataset (Figure 2), kallisto-bustools costs 39 times less than Salmon-Alevin-fry ($0.05 for kallisto-bustools vs. $1.96 for Salmon-Alevin-fry). This difference is substantial when pre-processing data for a genomics consortium, where there may be hundreds or thousands of datasets to analyze. The cost penalty for using Salmon-Alevin-fry will amount to thousands of dollars, and preclude routinely re-running the pre-processing datasets to update results with respect to updated annotations. Moreover, the cost establishes a barrier for other users to reproduce the results or to utilize the raw sequencing data. Even for individual research projects, which now regularly sample hundreds of thousands, or even millions of cells per experiment, the cost differential will be substantial and limiting. The high memory requirements of Salmon-Alevin-fry also preclude running the program on free cloud tools such as Google Colab, whereas kallisto-bustools is compatible with such infrastructure.

While one may suspect that the computational inefficiency of Salmon-Alevin-fry is due to the Alevin-fry reimplementation of the C++ bustools programs using the Rust programming language, we believe the differences are likely due to other reasons. One likely reason is Salmon-Alevin-fry’s use of “selective alignment with a decoy-aware transcriptome”. This alignment requires an indexing of the full genome alongside the transcriptome. The authors of Salmon-Alevin-fry have published several papers insisting that this practice improves results (Srivastava et al. 2020, 2021). Here we find that Salmon-Alevin-fry results on single-cell RNA-seq datasets using “selective alignment with a decoy-aware transcriptome” are nearly identical, insofar as downstream analyses are concerned, to those of kallisto, which performs pseudoalignment to a transcriptome only.

This raises the question of what the rationale would be for using Salmon-Alevin-fry, given that it provides no apparent advantage over kallisto-bustools, yet takes longer to run and uses more memory. We note that the Rust implementation of Alevin-fry does demonstrate that the Rust programming language holds promise for bioinformatics applications, and Rustaceans, who are increasingly common in the bioinformatics domain (Perkel 2020), may find Alevin-fry to be of pedagogical interest.

## Methods

The code to download the data, perform the analyses and obtain the results is located at https://github.com/pachterlab/BP_2021 and provides a complete description of the methods.

## Acknowledgements

We thank Diane Trout for sharing benchmarks that motivated this project. A.S.B. and L.P. were supported in part by NIH U19MH114830. The kallisto and bustools projects are supported in part by a grant for open source software from the Chan Zuckerberg Institute. We thank Mohsen Zakeri for suggesting in https://github.com/COMBINE-lab/BP_2021-lfl additions to the code-base to improve the reproducibility of the results in this manuscript.

