## Supplementary Material for "Benchmarking of lightweight-mapping based single-cell RNA-seq pre-processing"

### Supplementary Information

A. Sina Booeshaghi[1] and Lior Pachter[2]

[1] Department of Mechanical Engineering, California Institute of Technology
[2] Division of Biology and Biological Engineering and Department of Computing and Mathematical Sciences, California Institute of Technology

```
 1 $ cat logs/salmon_quant.log
 2 [2021-01-19 15:22:29.122] [jointLog] [info] setting maxHashResizeThreads to 10
 3 [2021-01-19 15:22:29.122] [jointLog] [info] Fragment incompatibility prior below threshold.
   Incompatible fragments will be ignored.
 4 [2021-01-19 15:22:29.122] [jointLog] [info] The --mimicBT2, --mimicStrictBT2 and --hardFilter flags
   imply mapping validation (--validateMappings). Enabling mapping validation.
 5 [2021-01-19 15:22:29.122] [jointLog] [info] Usage of --validateMappings implies use of
   minScoreFraction. Since not explicitly specified, it is being set to 0.65
 6 [2021-01-19 15:22:29.122] [jointLog] [info] The use of range-factorized equivalence classes does not
   make sense in conjunction with --hardFilter.  Disabling range-factorized equivalence classes.
 7 [2021-01-19 15:22:29.122] [jointLog] [info] Setting consensusSlack to selective-alignment default of
   0.35.
 8 [2021-01-19 15:22:29.122] [jointLog] [info] Using default value of 0.87 for minScoreFraction in Alevin
 9 Using default value of 0.6 for consensusSlack in Alevin
10 [2021-01-19 15:22:29.330] [jointLog] [info] There is 1 library.
11 [2021-01-19 15:22:29.415] [jointLog] [info] Loading pufferfish index
12 [2021-01-19 15:22:29.415] [jointLog] [info] Loading dense pufferfish index.
13 [2021-01-19 15:22:58.159] [jointLog] [info] done
14 [2021-01-19 15:22:58.159] [jointLog] [info] Index contained 110,411 targets
15 [2021-01-19 15:22:58.200] [jointLog] [info] Number of decoys : 66
16 [2021-01-19 15:22:58.200] [jointLog] [info] First decoy index : 110,310
17 [2021-01-19 16:02:37.043] [jointLog] [info] Computed 0 rich equivalence classes for further processing
18 [2021-01-19 16:02:37.043] [jointLog] [info] Counted 0 total reads in the equivalence classes
19 [2021-01-19 16:02:37.044] [jointLog] [info] Selectively-aligned 227833535 total fragments out of
   335147976
20 [2021-01-19 16:02:37.044] [jointLog] [info] Number of fragments discarded because they are best-mapped
   to decoys : 10,191,821
21 [2021-01-19 16:02:37.044] [jointLog] [warning] Found 1574 reads with `N` in the UMI sequence and
   ignored the reads.
22 Please report on github if this number is too large
23 [2021-01-19 16:02:37.044] [jointLog] [info] finished sc_align()
```

**Supplementary Figure 1:** Screenshot of the Salmon-Alevin-fry mouse-EMTAB7320_v2 salmon_quant.log file. Lines 7 describes setting the `consensusSlack` parameter to a default value of 0.35, and line 8 describes setting the same parameter to 0.6. Lines 5 and 8 describe setting the `minScoreFraction` to 0.65 and also to 0.87.

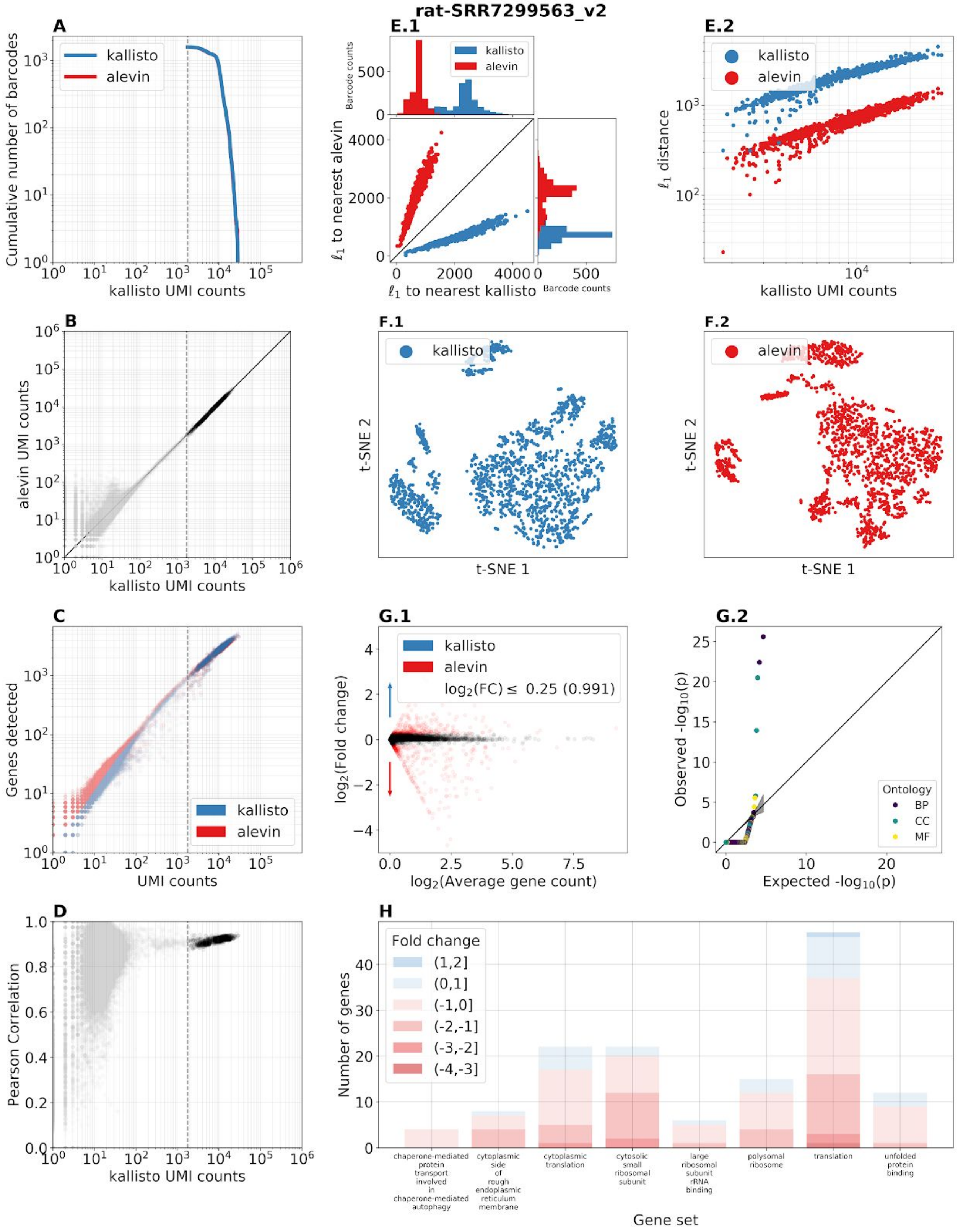


**Supplementary Figure 2.1:** Benchmark panel of dataset SRR7299563 from (Mays et al. 2018). [Code]

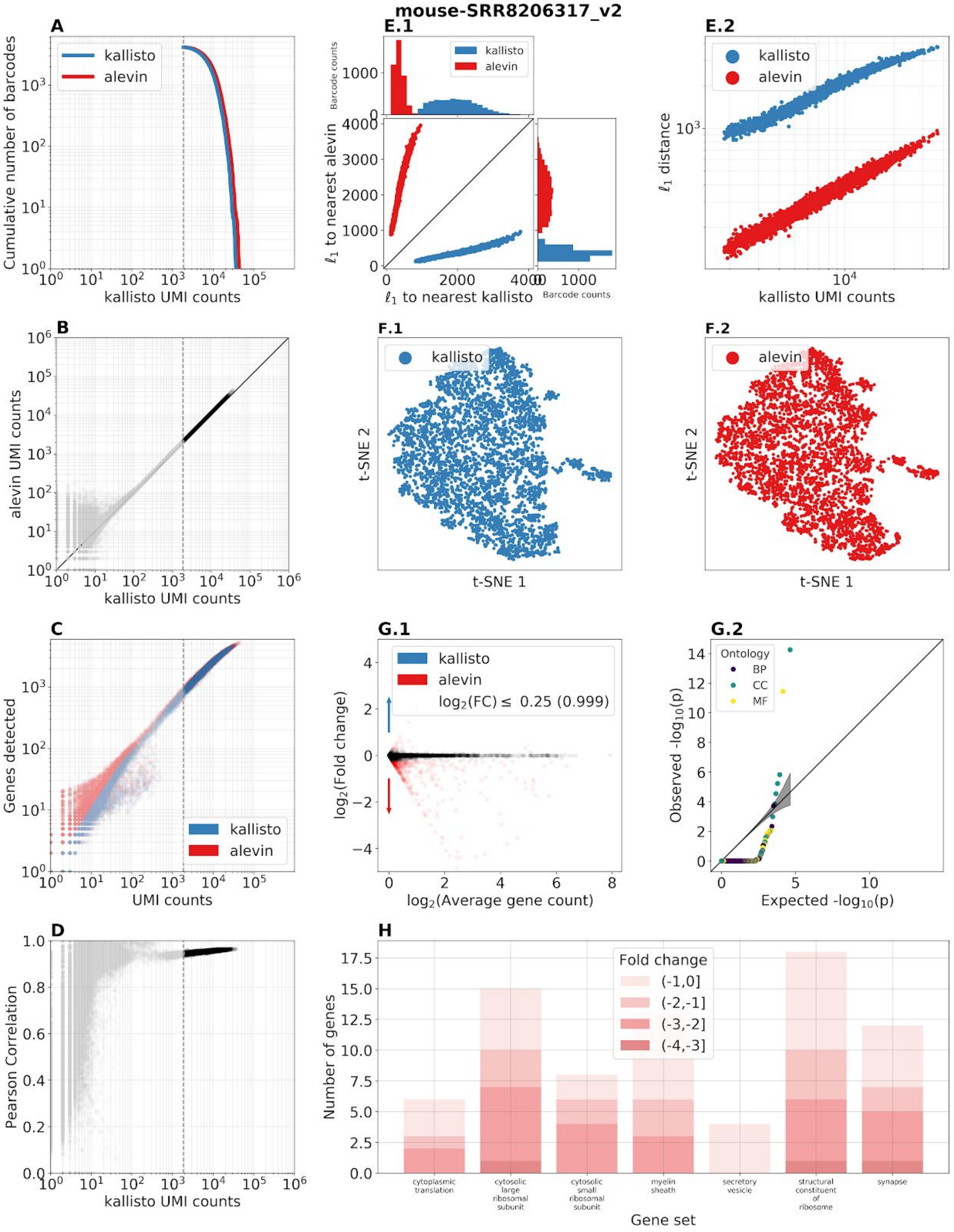


**Supplementary Figure 2.2:** Benchmark panel of dataset SRR8206317 from (Miller et al. 2019). [Code]

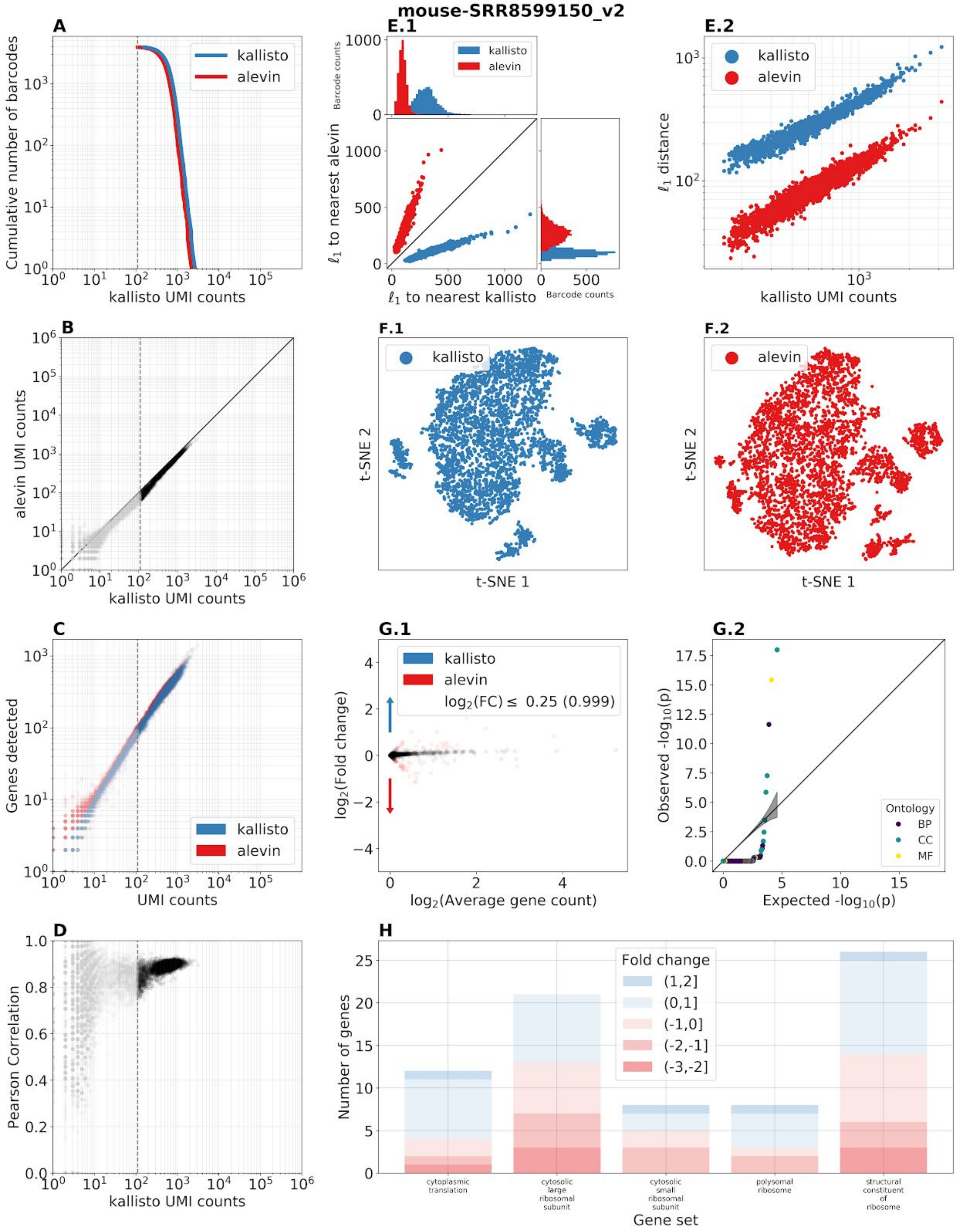


**Supplementary Figure 2.3:** Benchmark panel of dataset SRR8599150 from (O'Koren et al. 2019). [Code]

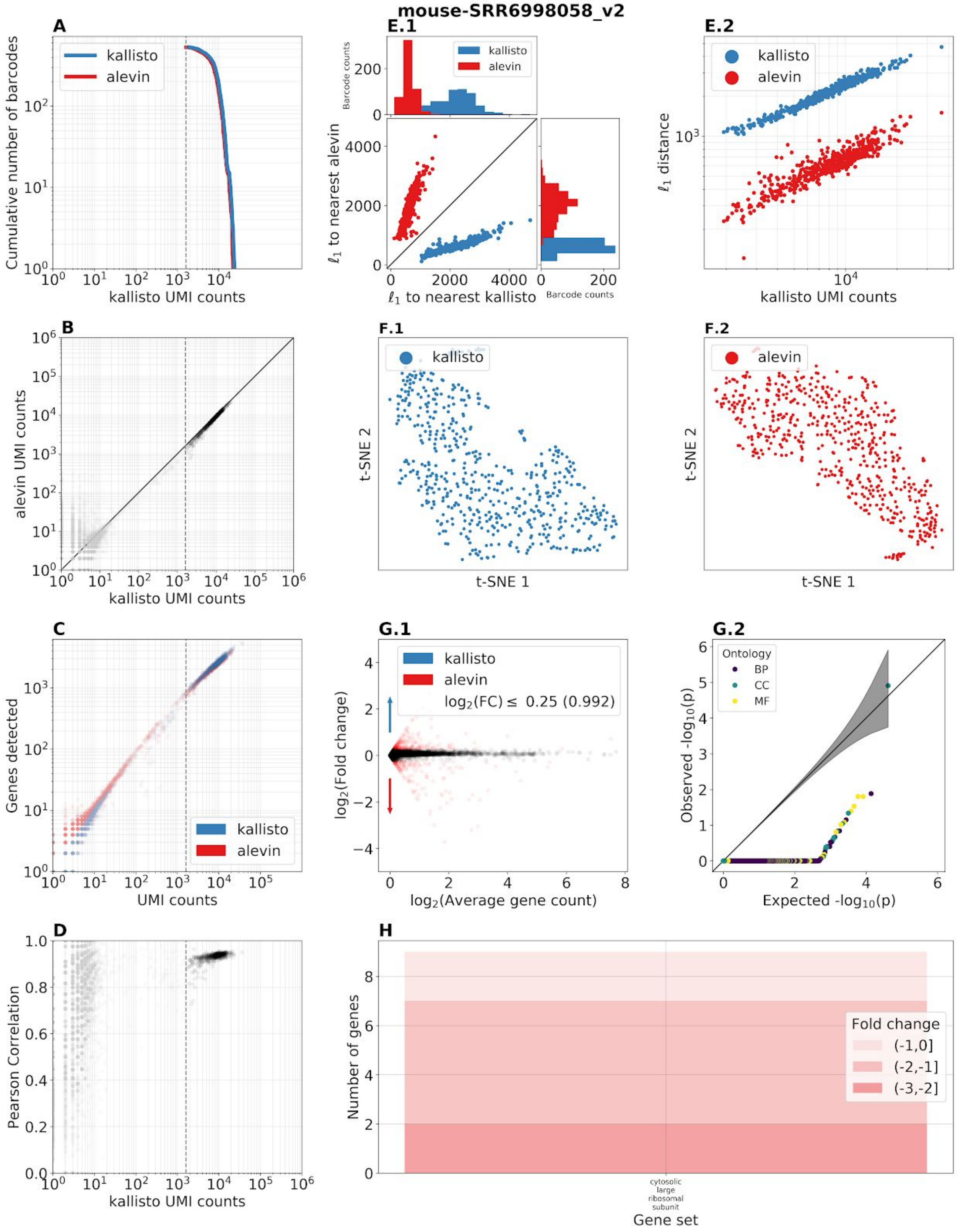


**Supplementary Figure 2.4:** Benchmark panel of dataset SRR6998058 from (Jin, Warunek, and Wohlfert 2018). [Code]

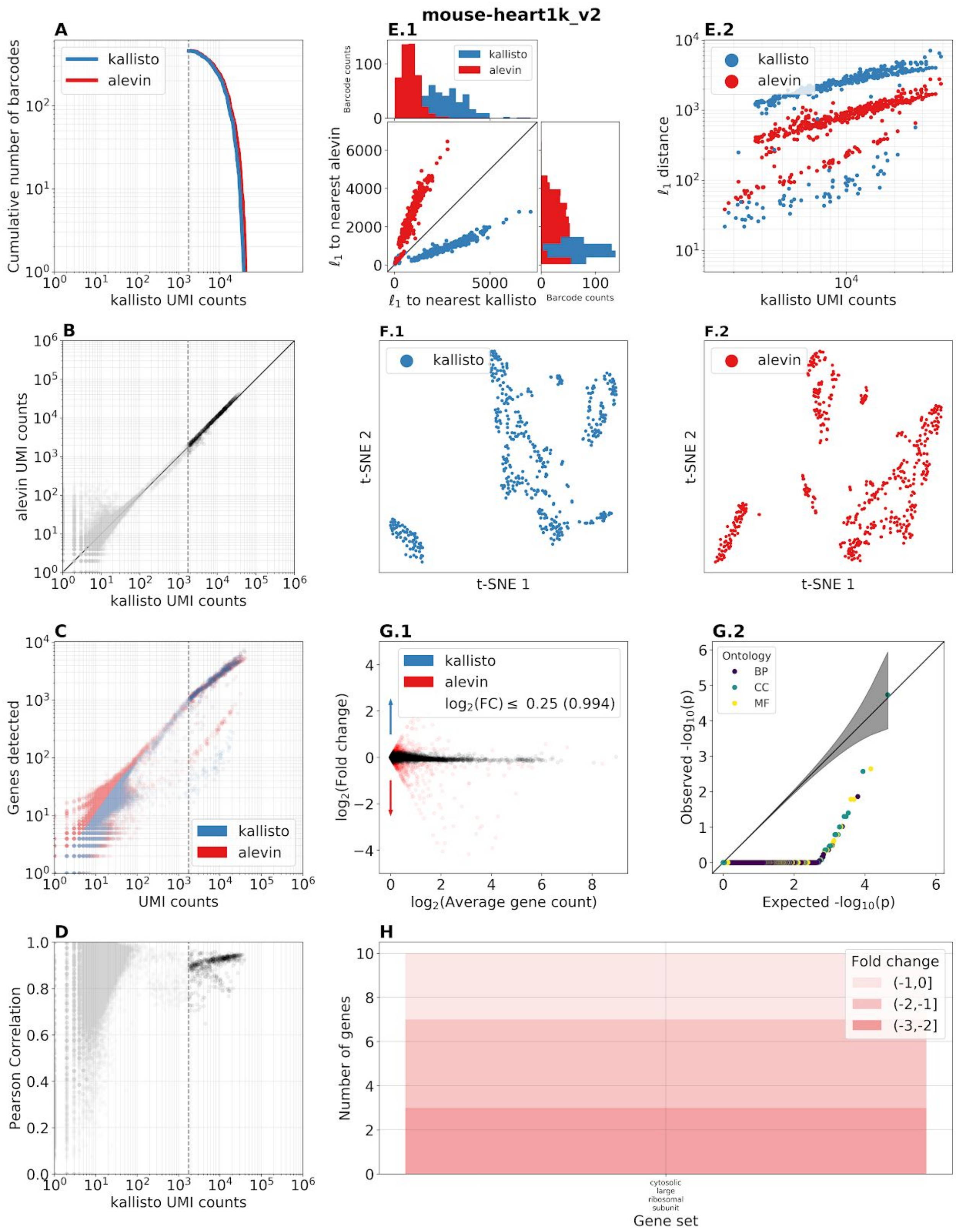


**Supplementary Figure 2.5:** Benchmark panel of the 10x Genomics heart1k_v2 dataset (“Datasets -Single Cell Gene Expression -Official 10x Genomics Support” n.d.). [Code]

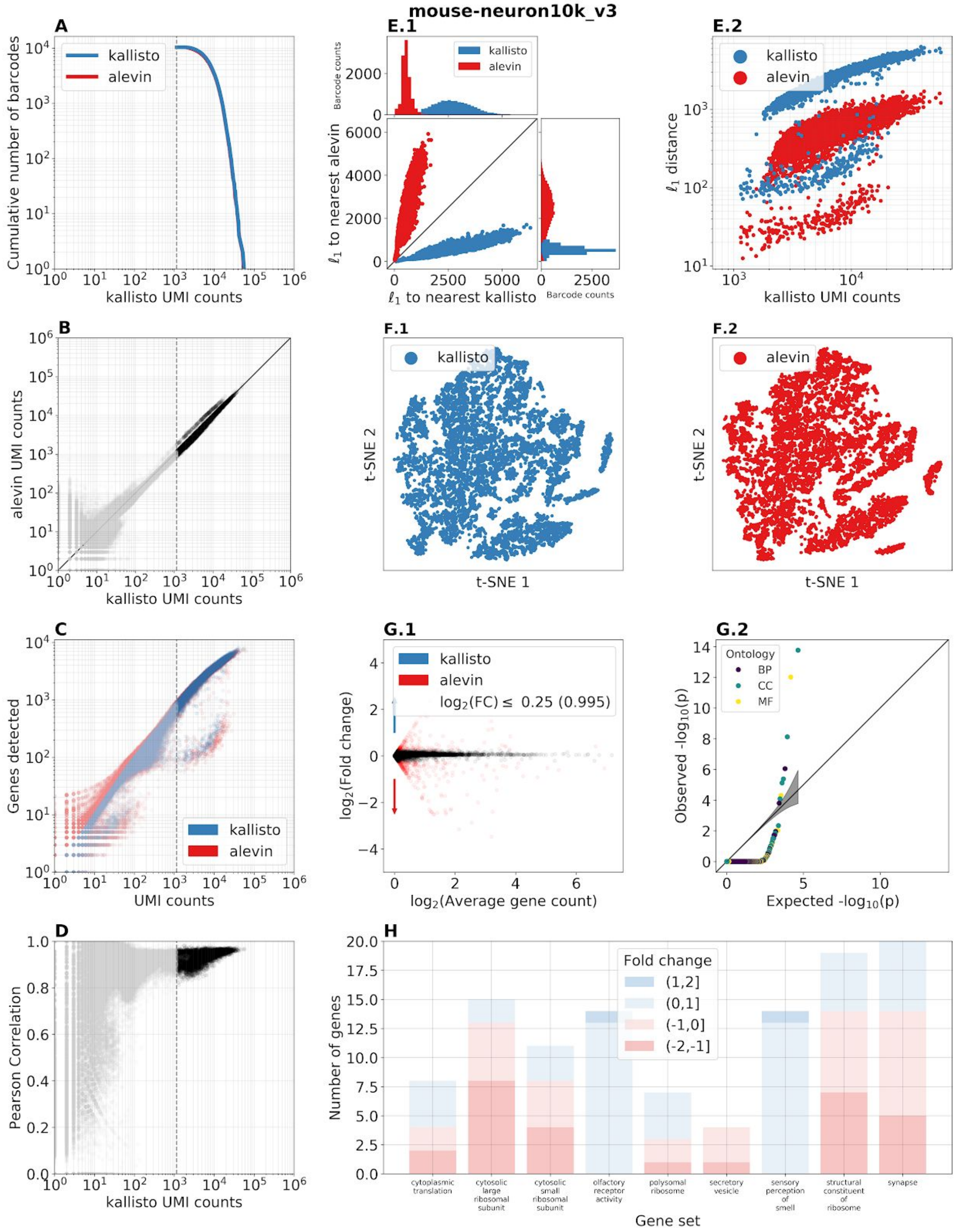


**Supplementary Figure 2.6:** Benchmark panel of the 10x Genomics neuron10k_v3 dataset (“Datasets -Single Cell Gene Expression -Official 10x Genomics Support” n.d.). [Code]

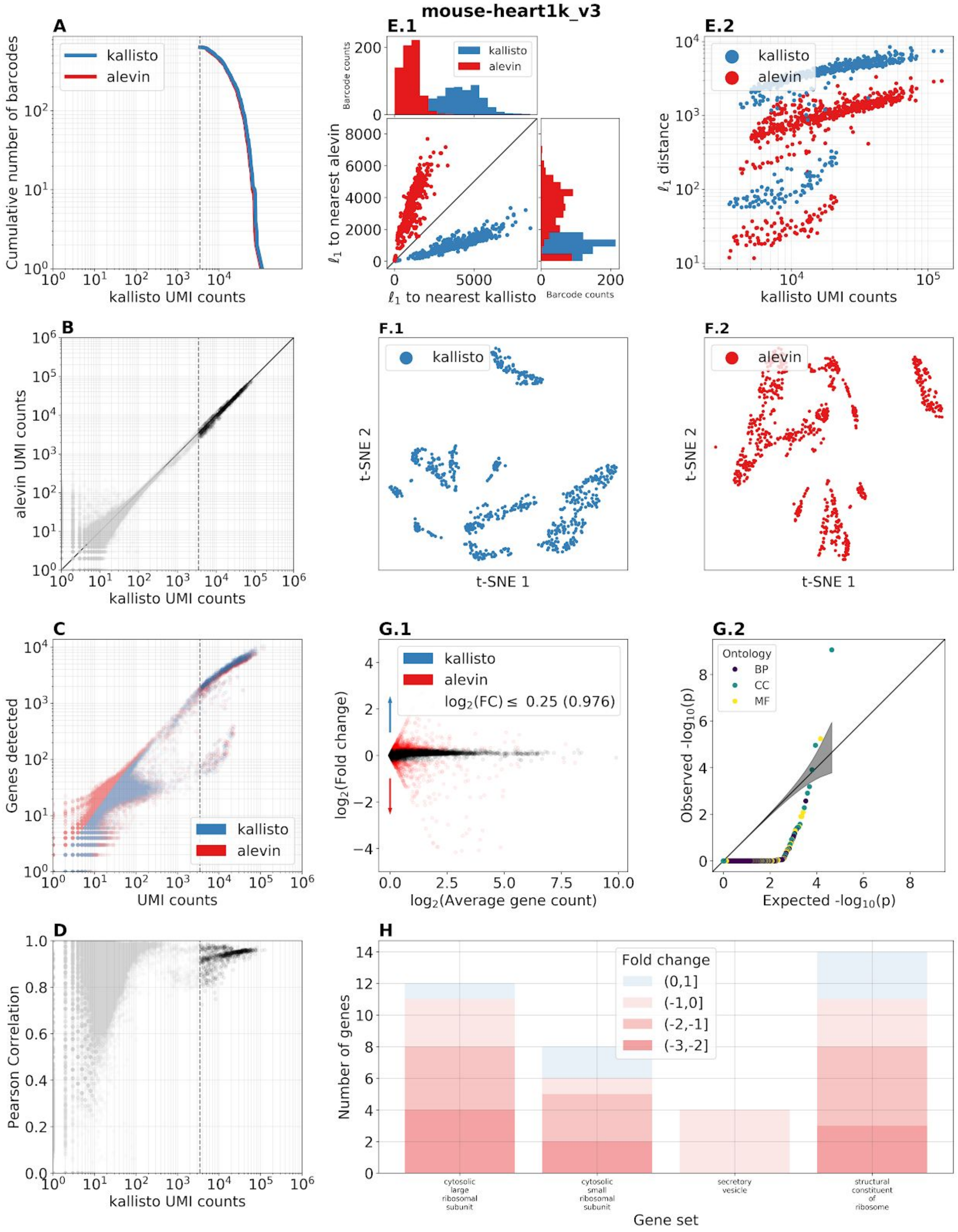


**Supplementary Figure 2.7:** Benchmark panel of the 10x Genomics heart1k_v3 dataset ("Datasets -Single Cell Gene Expression -Official 10x Genomics Support" n.d.). [Code]

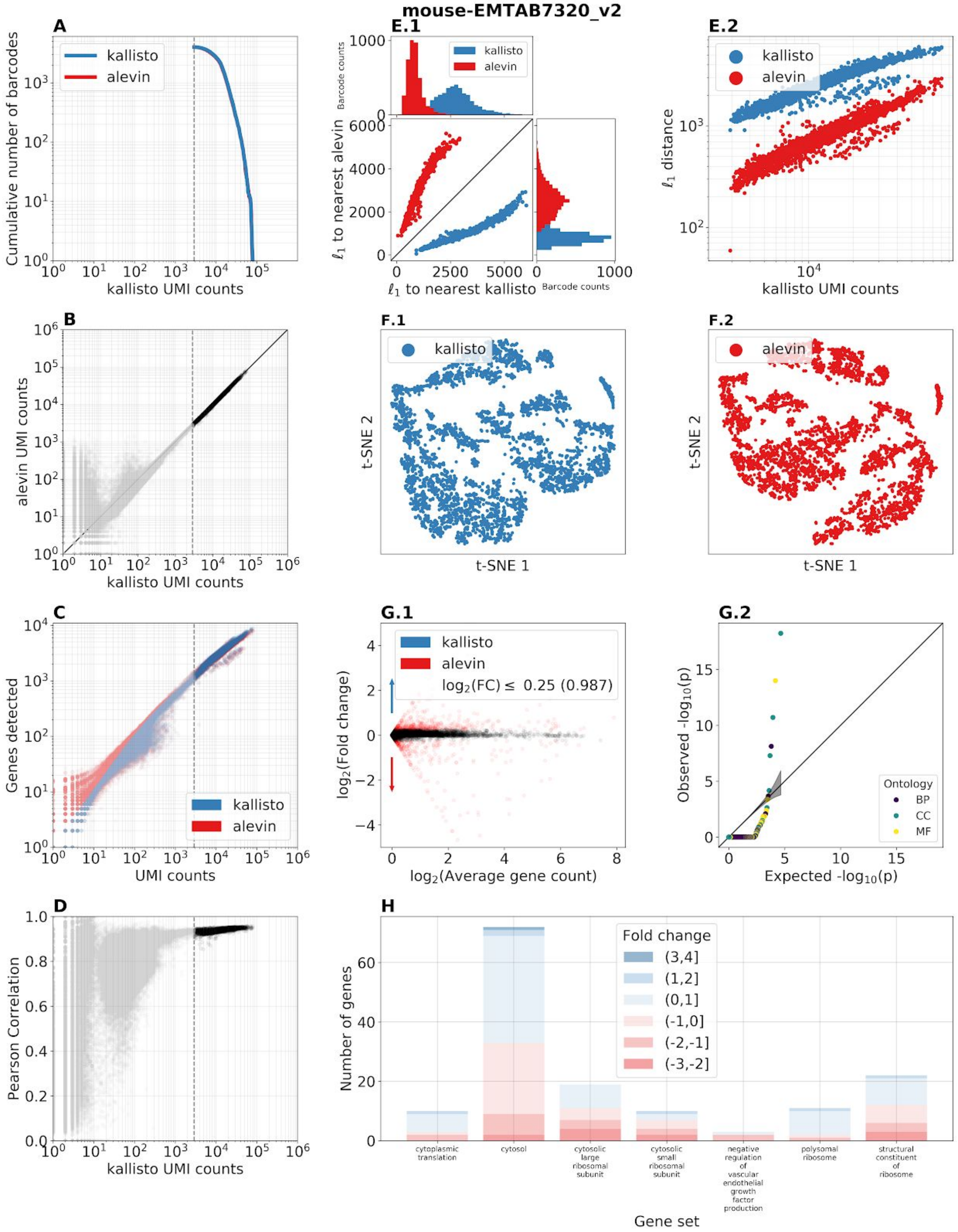


**Supplementary Figure 2.8:** Benchmark panel of dataset EMTAB7320 from (Delile et al. 2019). [Code]

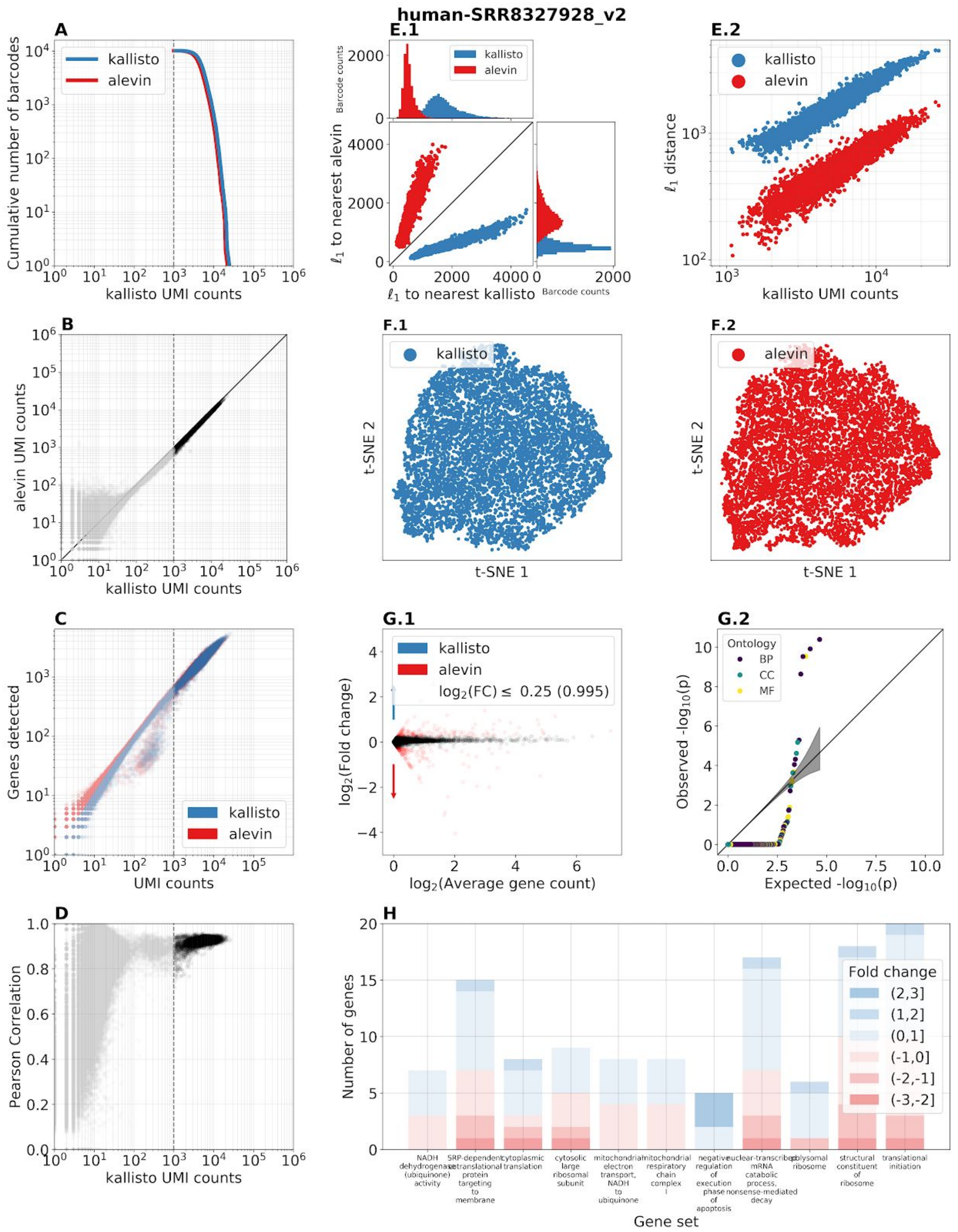


**Supplementary Figure 2.9:** Benchmark panel of dataset SRR8327928 from (Merino et al. 2019). [Code]

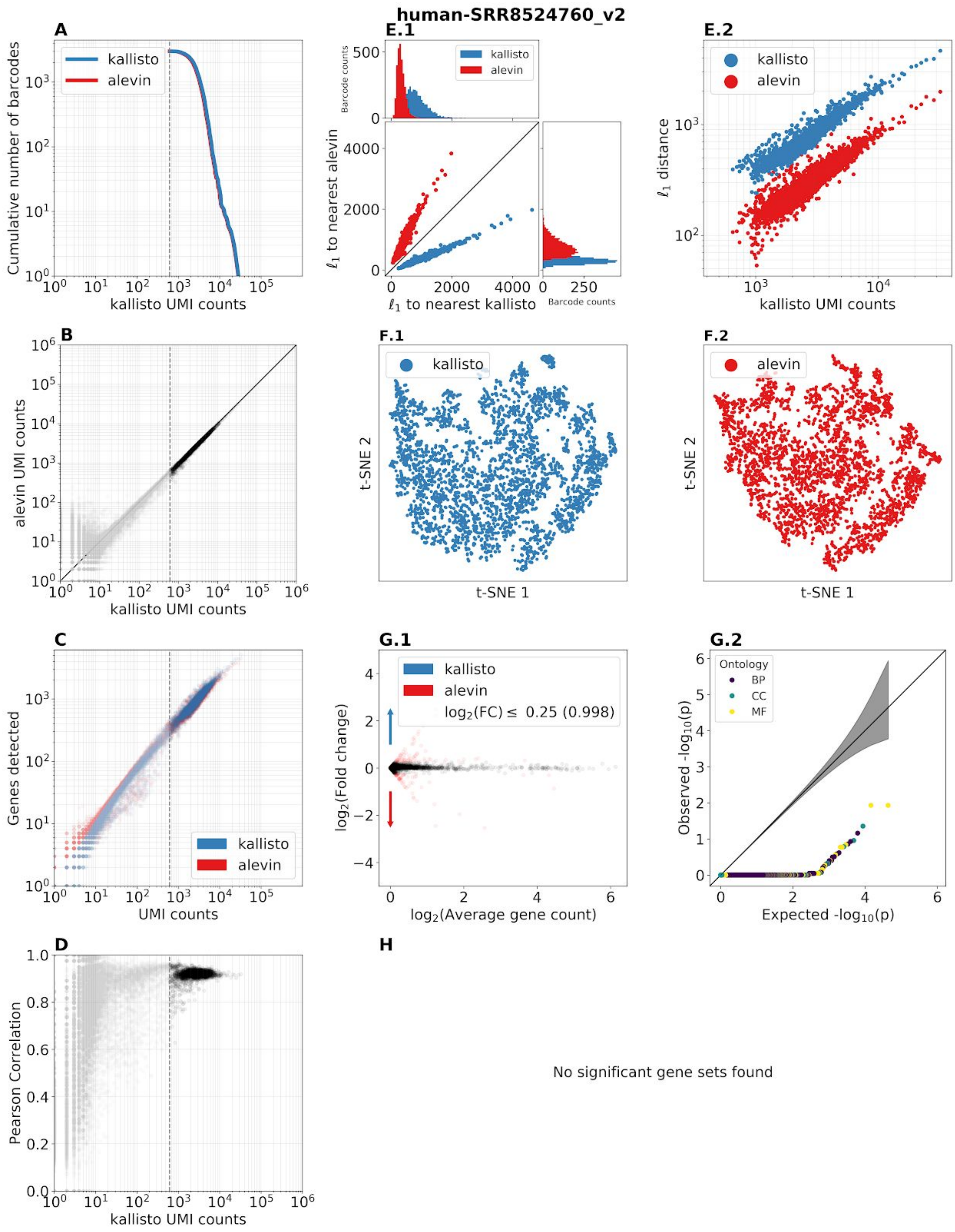


**Supplementary Figure 2.10:** Benchmark panel of dataset SRR8524760 from (Carosso et al. 2018). [Code]

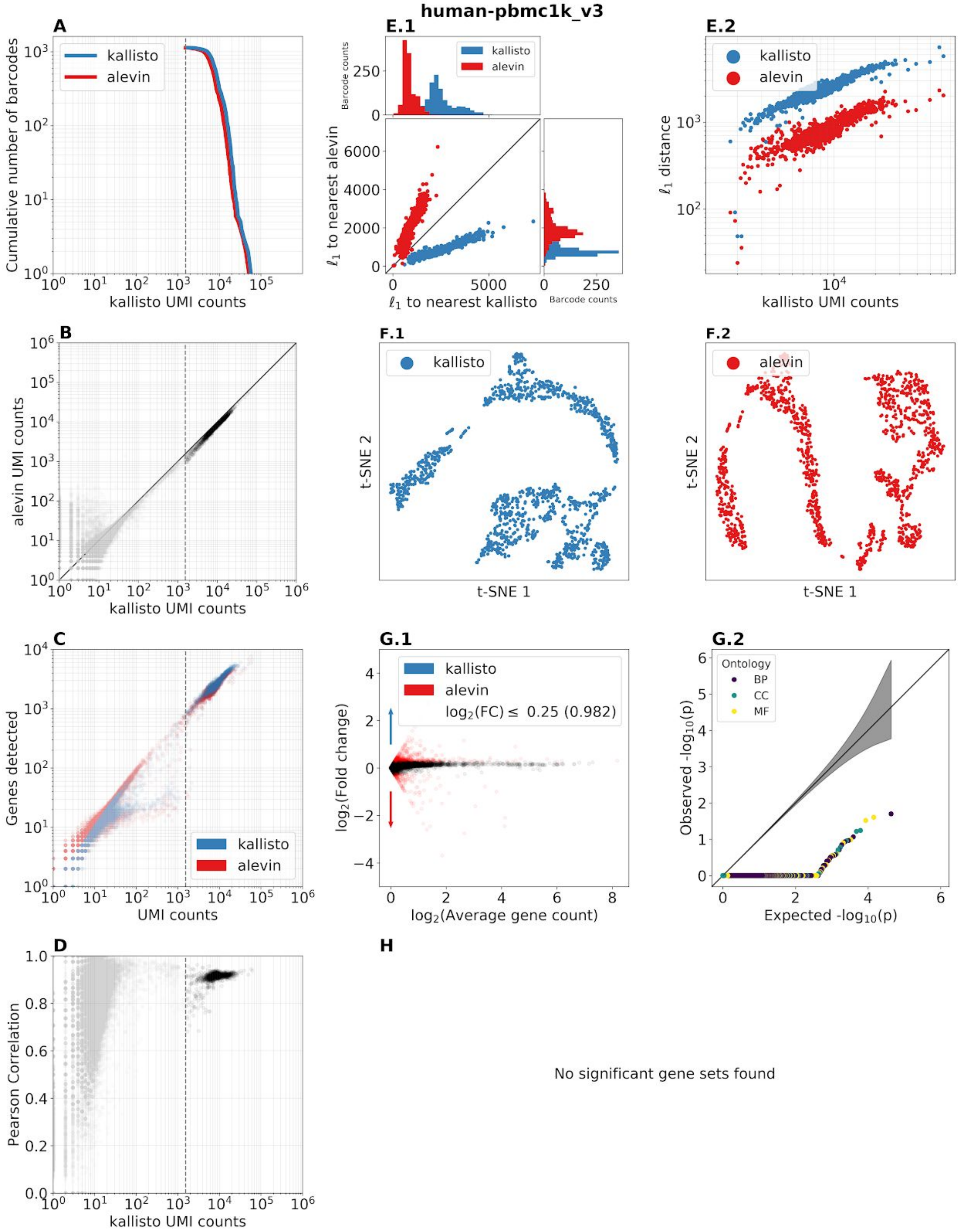


**Supplementary Figure 2.11:** Benchmark panel of the 10x Genomics pbmc1k_v3 dataset (“Datasets -Single Cell Gene Expression -Official 10x Genomics Support” n.d.). [Code]

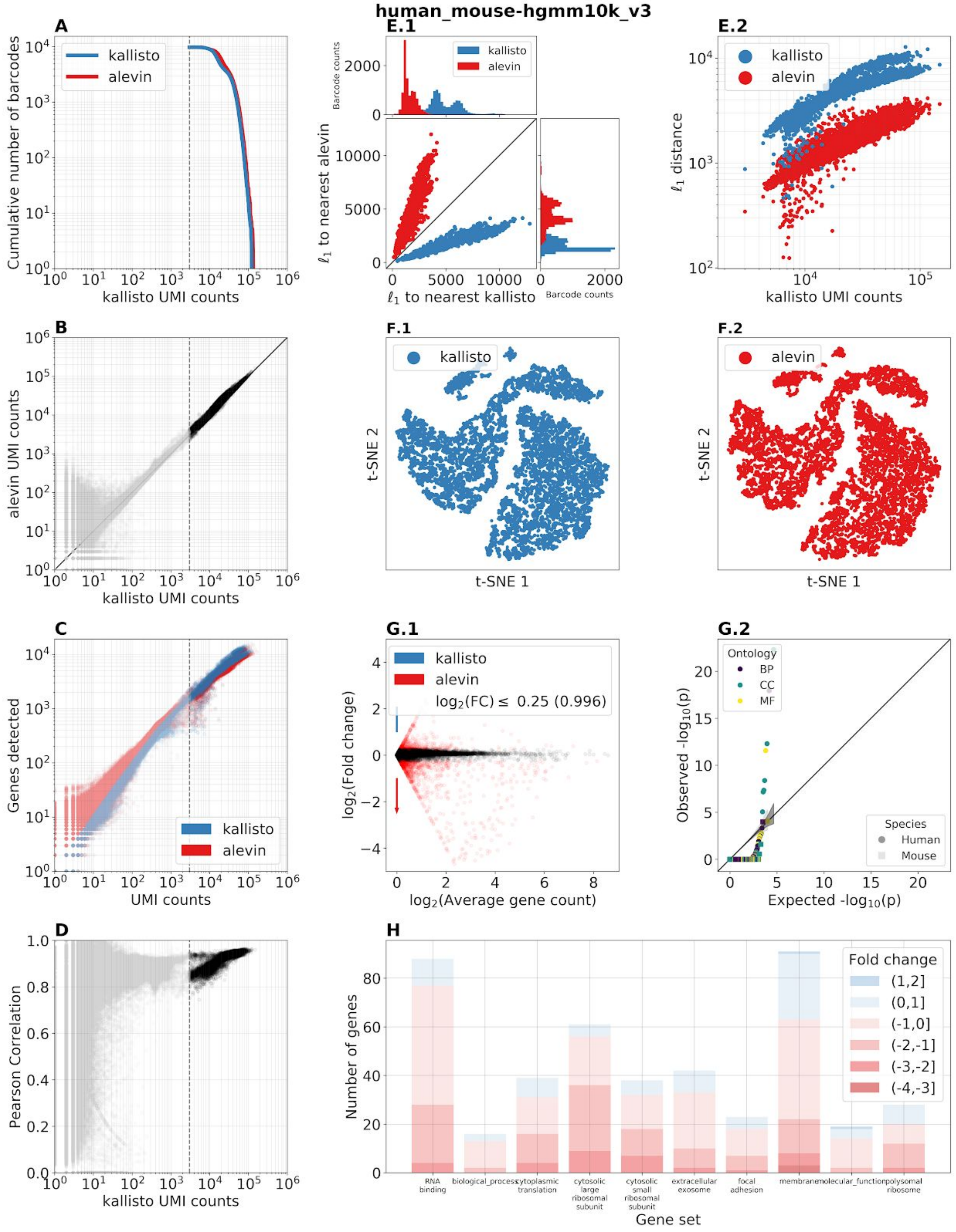


**Supplementary Figure 2.12:** Benchmark panel of the 10x Genomics hgmm10k_v3 dataset (“Datasets -Single Cell Gene Expression -Official 10x Genomics Support” n.d.). [Code]

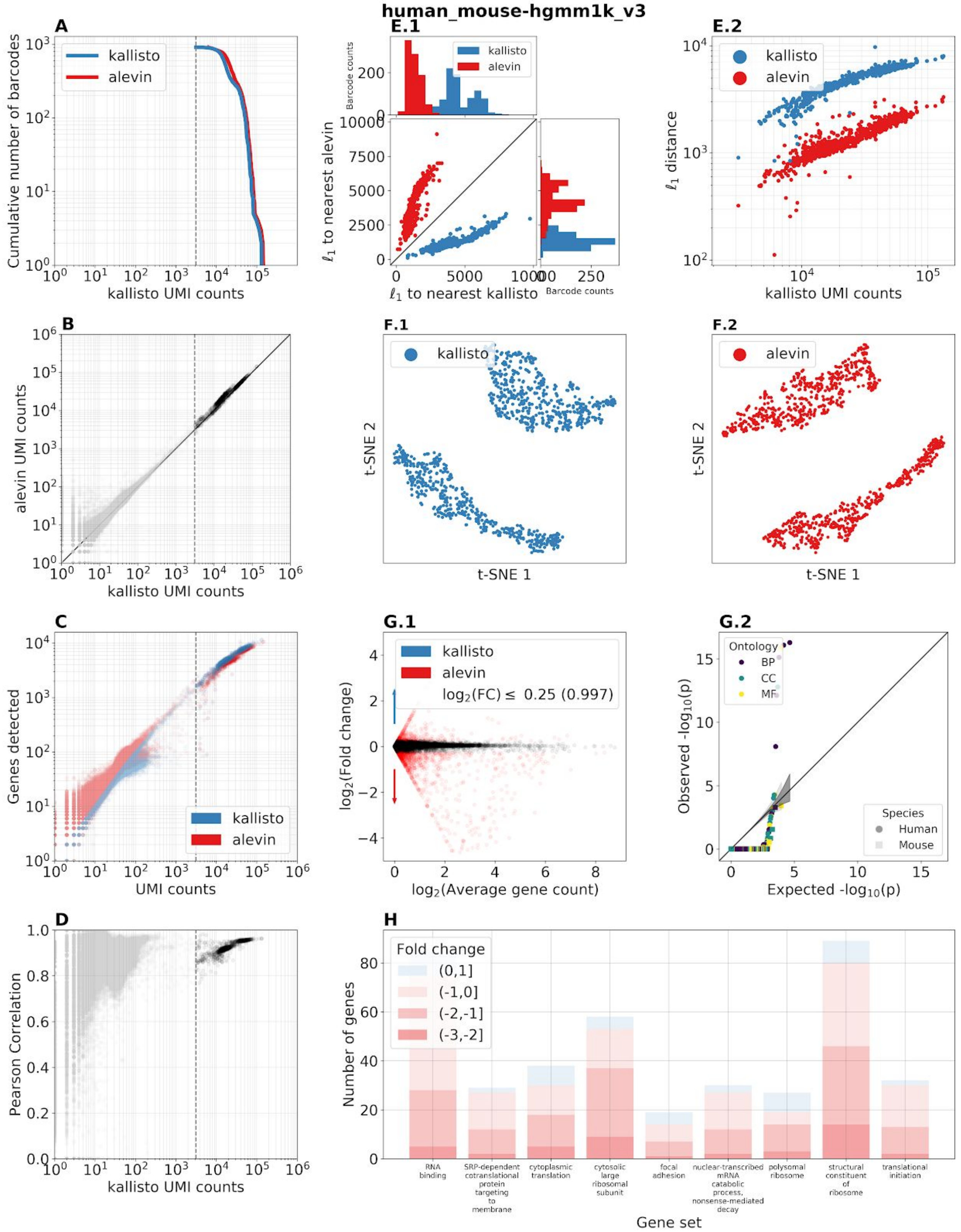


**Supplementary Figure 2.13:** Benchmark panel of the 10x Genomics hgmm1k_v3 dataset ("Datasets -Single Cell Gene Expression -Official 10x Genomics Support" n.d.). [Code]

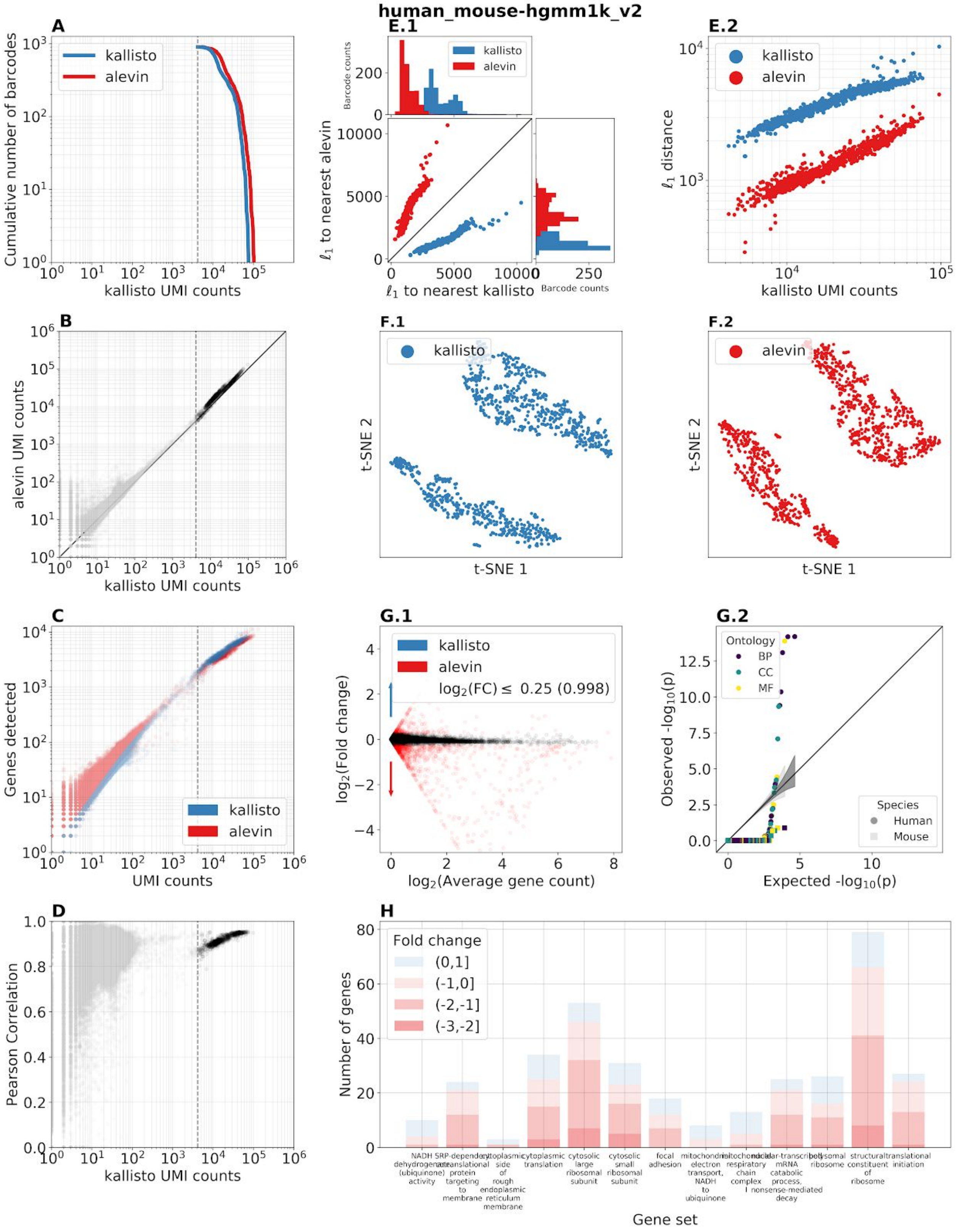


**Supplementary Figure 2.14:** Benchmark panel of the 10x Genomics hgmm1k_v2 dataset (“Datasets -Single Cell Gene Expression -Official 10x Genomics Support” n.d.). [Code]

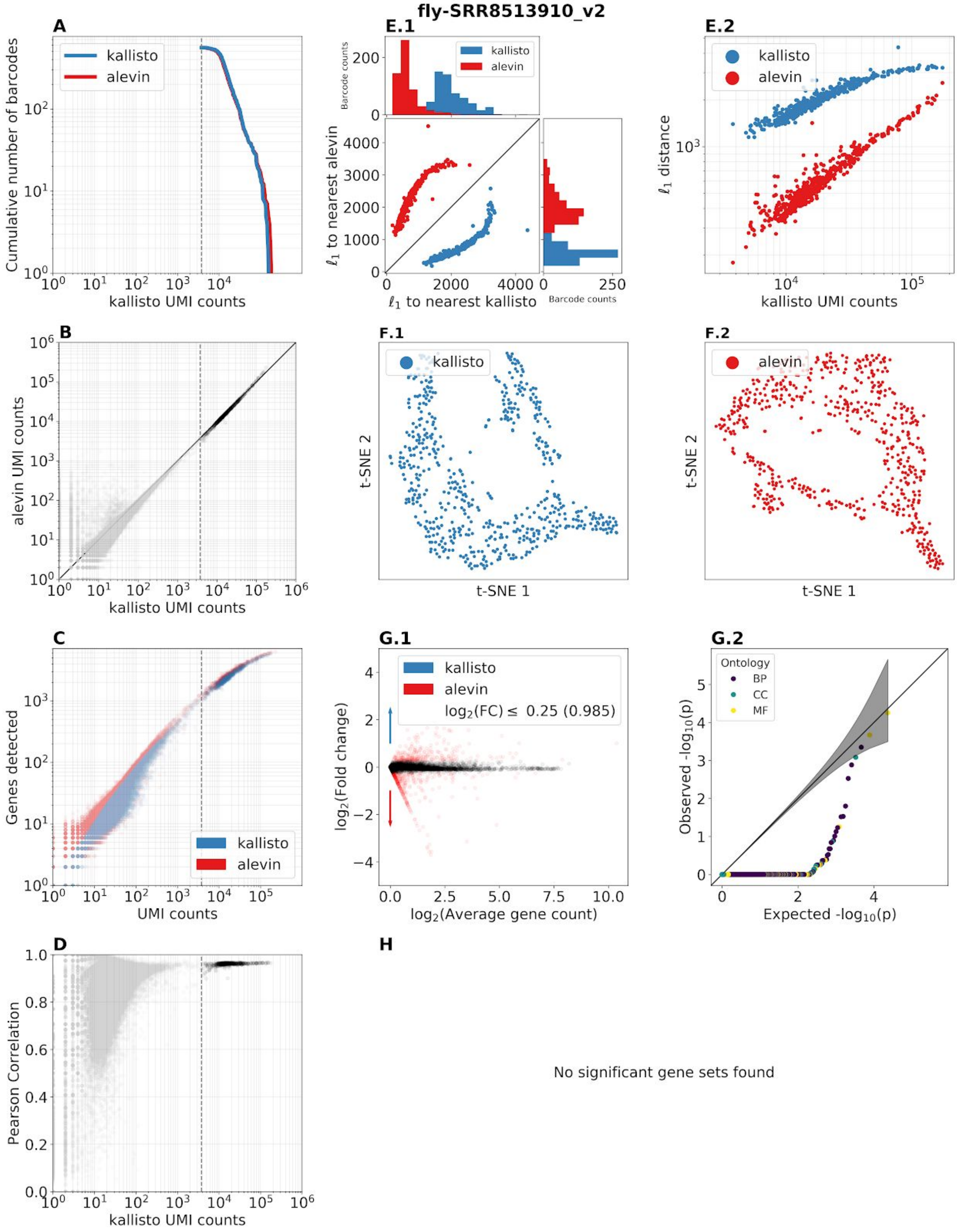


**Supplementary Figure 2.15:** Benchmark panel of dataset SRR8513910 from (Mahadevaraju et al. 2020). [Code]

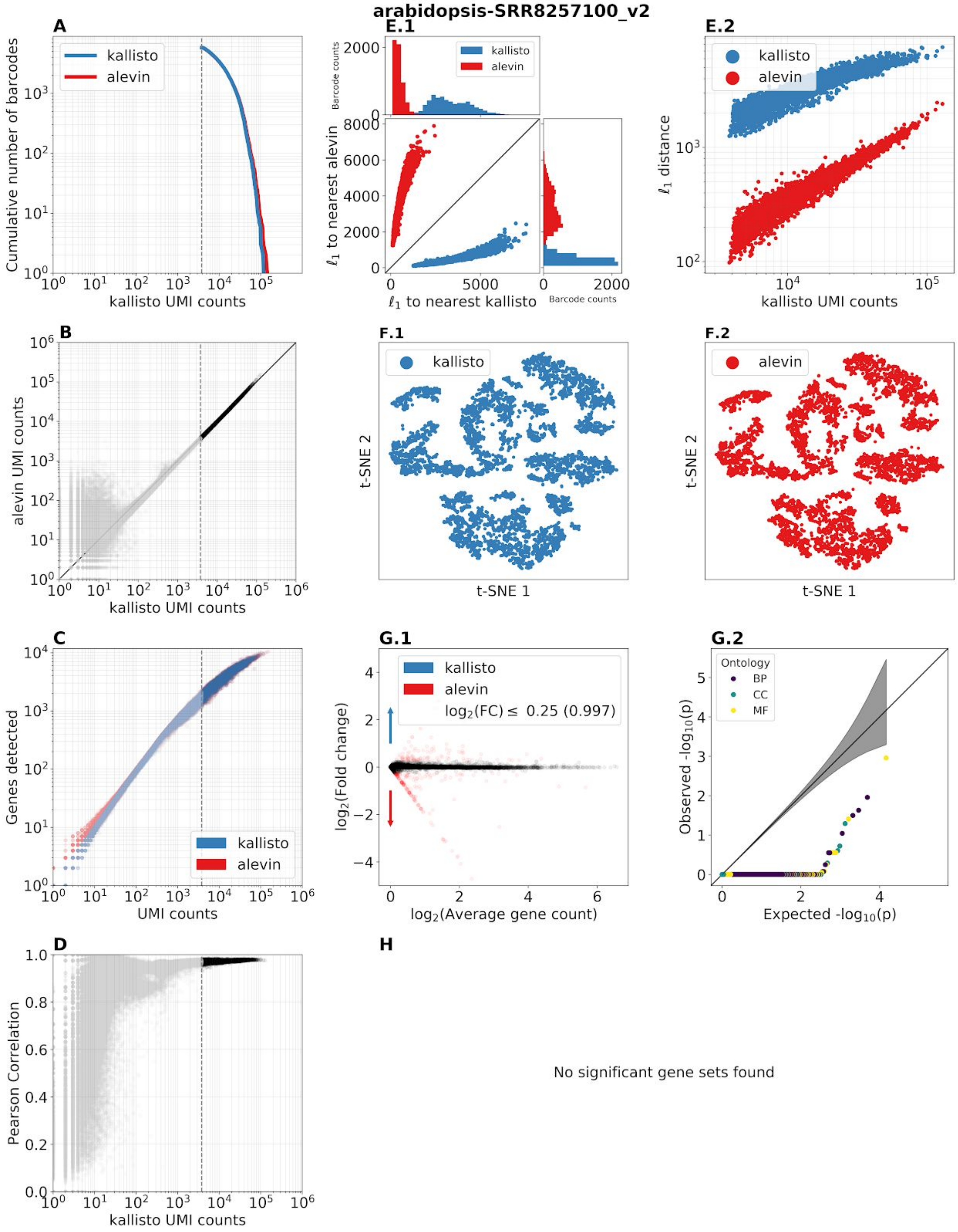


**Supplementary Figure 2.16:** Benchmark panel of dataset SRR8257100 (Ryu et al. 2019).
[Code]

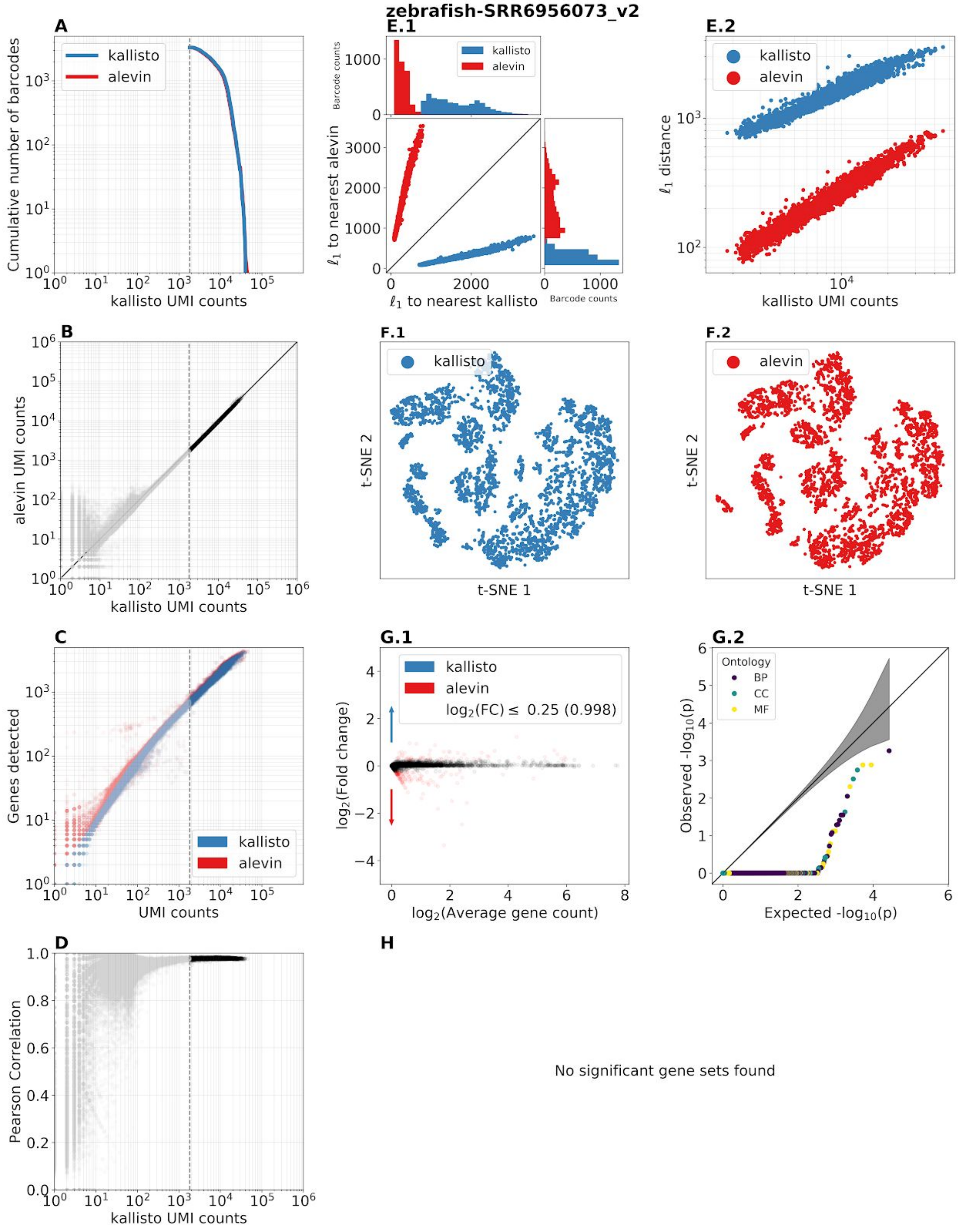


**Supplementary Figure 2.17:** Benchmark panel of dataset SRR6956073 from (Farrell et al. 2018). [Code]

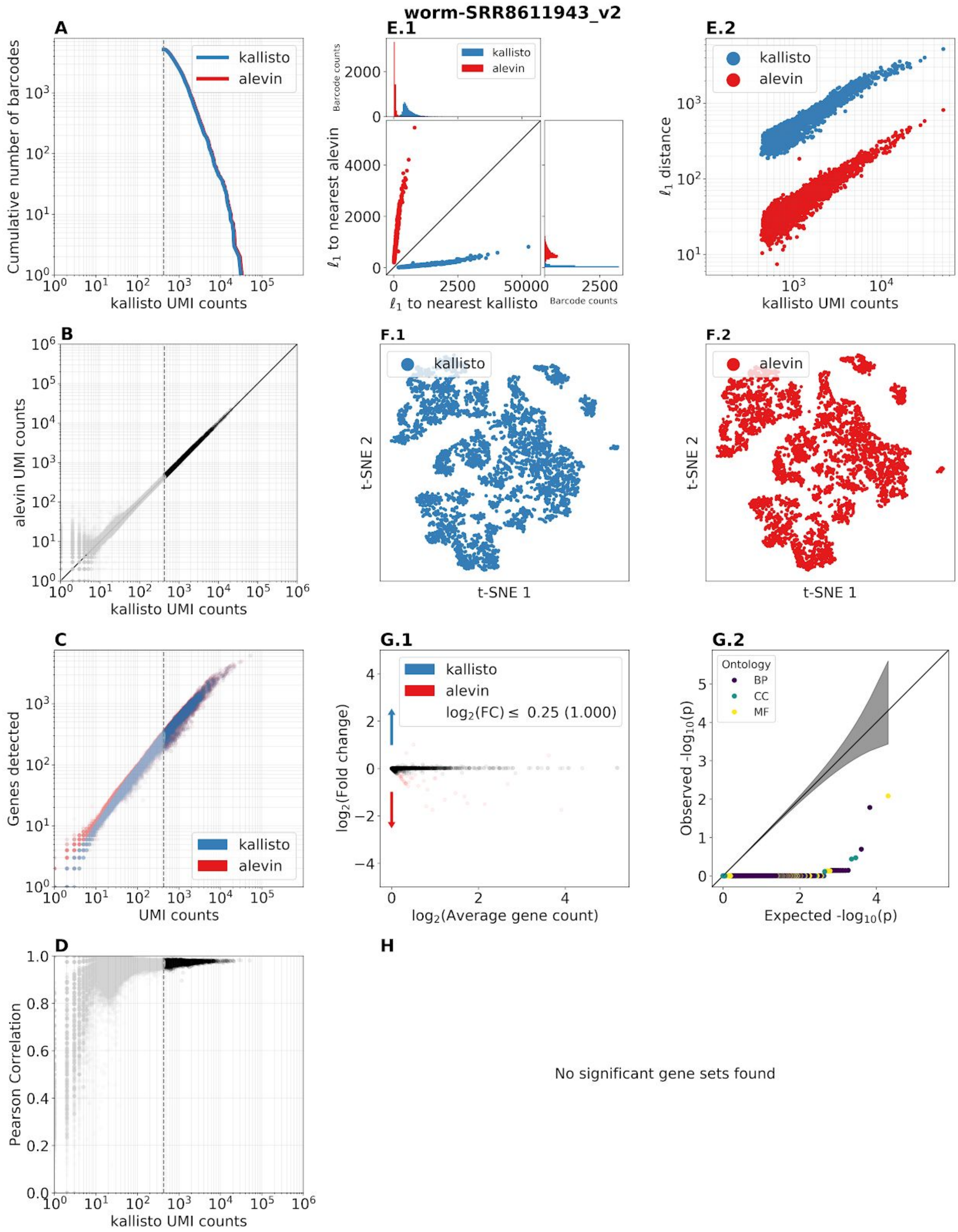


**Supplementary Figure 2.18:** Benchmark panel of dataset SRR8611943 from (Packer et al. 2019). [Code]

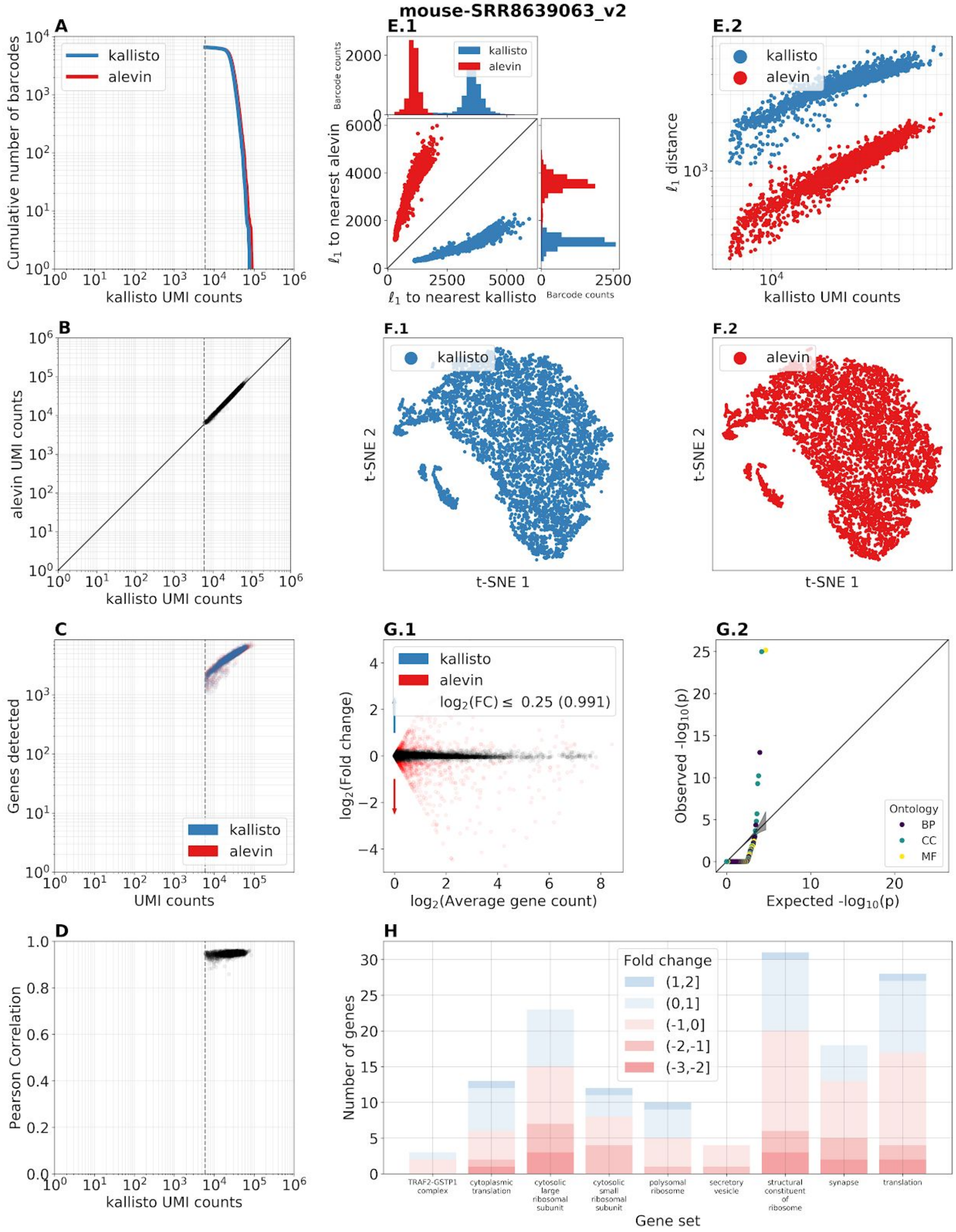


**Supplementary Figure 2.19:** Benchmark panel of dataset SRR8639063 from (Guo et al. 2019). The FASTQ files distributed with this experiment contained only filtered barcodes. [Code]

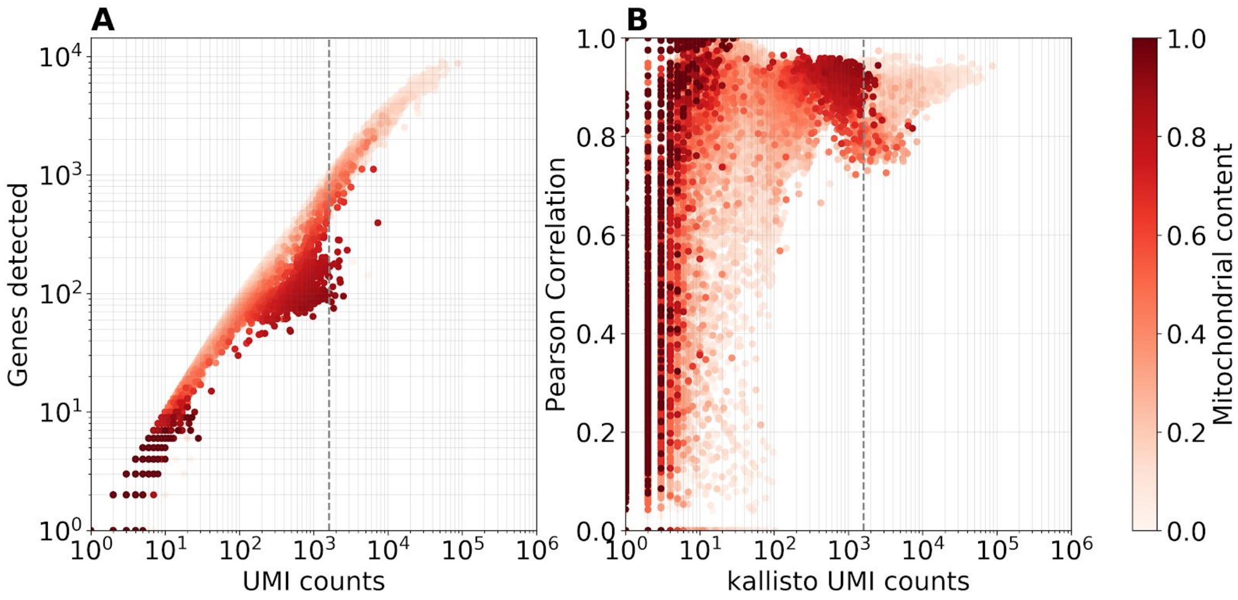


**Supplementary Figure 3:** Mitochondrial content of cells in the pbmc10k_v3 dataset. [Code]

| Workflow | kallisto-bustools | STARsolo | Salmon-Alevin-fry | RainDrop | Cell Ranger | Optimus |
|---|---|---|---|---|---|---|
| **Speed*** | 1x | 1.4x | 3.5x | - | 11.1x | 22.9x |
| **RAM** | 4 Gb | 64 Gb | 32 Gb | - | 32 Gb | 64 Gb |
| **Modular** | Yes | No | Yes | No | No | No |
| **Technology** | Multiple | Multiple | Multiple | 10xv2 only | 10x only | 10x only |
| **Assay** | Custom | scRNA-seq | Custom | 10x only | 10x only | scRNA-seq |
| **Mapping** | Lightweight | Alignment | Lightweight | Lightweight | Alignment | Alignment |
| **Ambiguity** | Yes | No | Yes | No | No | No |
| **Published** | Yes | No | Yes | Yes | No | No |
| **Depends** | C/C++ | C/C++ | C++/Rust | C++ | Python 2.7.13, Rust, C, Go, JavaScript | WDL, many programs |

**Supplementary Table 1:** Tools for pre-processing short-read single-cell RNA-seq data. The speed column benchmarks are derived from Supplementary Table 8 in *(Li et al. 2020). The RAM requirement is the minimum amount of RAM required by an AWS instance to run the associated tool. RainDrop was not benchmarked in Li et al. 2020. The “Modular” attribute refers to whether the code is modular, making it easy to swap out algorithms or design novel workflows. “Technology” refers to the single-cell RNA-seq technologies that are supported by the tool, “Assay” refers to the different types of workflows that can be designed, “Mapping” refers to the way in which reads are assigned to a reference, “Ambiguity” refers to whether the program assigns reads that are ambiguous to transcripts, “Published” refers to whether the workflow is published, and “Depends” refers to the dependencies required to run the program.

Ryu, Kook Hui, Ling Huang, Hyun Min Kang, and John Schiefelbein. 2019. “Single-Cell RNA Sequencing Resolves Molecular Relationships Among Individual Plant Cells.” *Plant*

*Physiology* 179 (4): 1444–56.
